# Interferon-gamma Induces Melanogenesis Via Post-Translational Modification of Tyrosinase

**DOI:** 10.1101/2021.10.14.464471

**Authors:** Xuan Mo, Hasan Raza Kazmi, Sarah Preston-Alp, Bo Zhou, M. Raza Zaidi

## Abstract

Melanogenesis (melanin pigment production) in melanocytes is canonically stimulated by the alpha-melanocyte stimulating hormone (αMSH), which activates the cyclic-AMP-mediated expression of the melanocyte inducing transcription factor (MITF) and its downstream melanogenic genes, including the principal rate-limiting melanogenic enzyme tyrosinase (Tyr). Here we report that interferon-gamma (IFNG; type II interferon), but not IFN-alpha (a type I interferon), induces a noncanonical melanogenic pathway. Inhibition of IFNG pathway by the JAK inhibitor ruxolitinib or knocking out Stat1 abrogated the IFNG-induced melanogenesis. Interestingly, IFNG-induced melanogenesis was independent of MITF. IFNG markedly increased the Tyr protein expression but did not affect the mRNA expression, suggesting a post-translational regulatory mechanism. In contrast, IFNG had no effect on the expression of other melanogenesis-related proteins, e.g. tyrosinase-related protein 1 (Tyrp1) and dopachrome tautomerase (Dct). Glycosidase digestion assays revealed that IFNG treatment increased the mature glycosylated form of Tyr, but not its *de novo* synthesis. Moreover, cycloheximide chase assay showed that degradation of Tyr was decreased in IFNG-treated cells. These results suggest that the IFNG-STAT1 pathway regulates melanogenesis via modulation of the post-translational processing and protein stability of Tyr.

**SIGNIFICANCE:** The canonical pathway that controls melanogenesis in melanocytes is activated by the alpha melanocyte stimulating hormone (αMSH) via its receptor MC1R, which activates the cyclic-AMP-mediated expression of the melanocyte master regulator MITF and its downstream target genes involved in the melanogenic process. Here we report a novel non-canonical melanogenic pathway that is mediated via the Interferon-gamma cytokine signaling. We show that this non-canonical pathway is independent of MITF-mediated gene expression, but rather functions via post-translational modification of the principal melanogenic enzyme Tyrosinase.

## 1. INTRODUCTION

Melanin is the principal contributor to the mammalian skin and hair pigmentation. It is synthesized in melanosomes, which are lysosome-related cellular organelles within melanocytes. Melanosomes are transported to the adjacent keratinocytes leading to skin pigmentation, which serves as a key physiological defense against the genotoxic effects of ultraviolet radiation (UVR) (Mo et al., 2019). Inflammatory cytokines secreted by keratinocytes and infiltrated lymphocytes in the skin microenvironment after UV irradiation or other dermatological damages have been implicated in regulating proliferation, differentiation, and melanogenesis in melanocytes via direct or indirect mechanisms (Choi et al., 2013; Wang et al., 2013). The most well-studied and potent inducer of melanogenesis is alpha-melanocyte-stimulating hormone (αMSH), which is synthesized in keratinocytes following exposure to UVR, secreted into the skin microenvironment, and stimulates melanogenesis in melanocytes. αMSH activates melanocyte-inducing transcription factor (MITF), which is considered to be the principal regulator of melanin biosynthesis (Nguyen and Fisher, 2019). MITF initiates transcription of numerous genes coding for melanogenesis-associated proteins, including tyrosinase (TYR), tyrosinase-related protein 1 (TYRP1), and Dopachrome tautomerase (DCT) (Kawakami and Fisher, 2017; Sturm, 2006). Tyrosinase catalyzes the tyrosine oxidation to dopaquinone, which is the first and rate-limiting step of melanogenesis (Chang, 2009). TYRP1 and DCT are involved in later steps of melanin synthesis and also play important roles in regulating the melanogenic apparatus (Slominski et al., 2004).

Tyrosinase is a type I membrane-bound glycoprotein (Jagirdar et al., 2014). The nascent TYR protein is cotranslationally translocated into the lumen of the endoplasmic reticulum (ER) and undergoes initial asparagine-linked (N-linked) glycosylation (Negroiu et al., 1999). The essentiality of glycosylation to the enzymatic function of TYR has been demonstrated, as inhibition of glycosylation with tunicamycin or glucosamine abolishes the maturation, trafficking, and the enzymatic activity of TYR (Halaban et al., 1997; Wang et al., 2006). Furthermore, mutations of the TYR N-linked glycosylation sites cause hypopigmentation phenotype by obstructing tyrosinase maturation (Branza-Nichita et al., 2000). The core-glycosylated TYR (∼70kD) is transported to the trans-Golgi network (TGN), where it undergoes further modifications of additional mannose residues after passing the quality control in the ER. Mature TYR (∼ 80kD) is then transported out of the TGN to the melanosomes where melanin synthesis occurs (Wang et al., 2006). Aberrant disruption of tyrosinase trafficking also affects melanin production. Regulation of melanogenesis via alteration of tyrosinase occurs at both transcriptional and post-transcriptional levels.

Interferon-gamma (IFNG) has long been known as the principal proinflammatory cytokine that plays the role of the central orchestrator of inflammation and autoimmune disease (Schroder et al., 2004). IFNG is predominantly produced by the NK cells, NK-T cells, and macrophages of the innate immune system, as well as the CD4+ Th1 and CD8+ cytotoxic T lymphocytes of the adaptive immune system (Schroder et al., 2004; Zaidi, 2019; Zaidi and Merlino, 2011). IFNG is directly involved in almost all types of skin inflammatory conditions, e.g. those caused by pathogenic infections, tissue injury, tissue stress, and/or malfunction (Dries and Perry, 2002; Xiao et al., 2009). IFNG signaling pathway is a well-established regulator of the classical activation of macrophages, and thus controls the synthesis and secretion of cytokines and enzymes important for tissue remodeling and wound healing (Fujiwara and Kobayashi, 2005; Schroder et al., 2004).

UVR induces an inflammatory response in the skin in which IFNG plays an important orchestrating role. IFNG receptors, IFNGR1 and IFNGR2, transduce IFNG signaling from the extracellular environment to the cellular signaling machinery. The IFNGR1 is associated with one member of the Janus activated kinase family, JAK1, whereas IFNGR2 is associated with JAK2 (Avalle et al., 2012). The canonical IFNG signaling is mediated by JAK/STAT (signal transducer and activator of transcription) pathway. The receptors are rearranged and dimerized upon binding of IFNG, followed by the autophosphorylation and activation of receptor-associated JAK1 and JAK2. The activated JAKs phosphorylate the tyrosine residue at position 701 (Tyr701) in STAT1, leading to the formation and activation of STAT1 homodimers (Wenta et al., 2008), which translocate to the nucleus and bind to the IFNG-activated site (GAS) elements on their target loci and initiate transcription of IFNG-stimulated genes to mediate numerous biological responses (Zaidi, 2019). Here we present evidence that IFNG signaling pathway is a regulator of melanogenesis via post-translational modification of TYR.

## 2. MATERIALS AND METHODS

### 2.1 Cell Culture

Mouse melanoma cell lines B16, B16N (a subclone of B16 cell line, generated at the National Cancer Institute, NIH), and human melanoma cell line Hs 936.T (ATCC) were cultured in Dulbecco’s Modified Eagle Medium (DMEM) supplemented with 10% fetal bovine serum (FBS), L-alanyl-L-Glutamine (2 mM) and Gentamycin (50 ug/ml). Cells were grown at 37 °C in a humidified incubator under 5% CO2.

### 2.2 Cytokine Treatment

Mouse recombinant IFN-gamma (Ifng) was purchased from Cell Signaling Technology (catalog #5222). The typical concentration used was 10 ng/ml, unless otherwise indicated. Mouse IFN-alpha 2 (Ifna2) recombinant protein was purchased from Affymetrix eBioscience (catalog #14-8312). The typical concentration used was 10 ng/ml, unless otherwise indicated. Alpha melanocyte-stimulating hormone (αMSH) was purchased from Sigma-Aldrich (catalog #581-05-5). The concentration used for mouse melanoma cells and human melanoma cells were 10 nM and 100 nM, respectively. Human recombinant IFN-gamma (IFNG) was purchased from Cell Signaling Technology (catalog #8901). The concentration used was 100 ng/ml. Media and cytokines were refreshed every other day.

### 2.3 Melanin Content Assay

Melanoma cells were collected after the treatment with different cytokines for indicated time periods. The pellets were lysed in NP-40 lysis buffer containing 1x protease inhibitors. Melanin was separated from lysates by centrifugation for 15 min, then dissolved in 1 M NaOH for 1 h at 100º C. The melanin content was determined by measuring the absorbance at 490 nm using a microplate reader. The synthetic melanin standard (Sigma) curve was generated to calculate the melanin content of each sample. The protein concentration of each sample was measured with the Bio-Rad Protein Assay following the manufacturer’s protocol. The melanin content was obtained by normalization of melanin amount to protein input.

### 2.4 Ruxolitinib Treatment

B16N cells were pretreated with or without ruxolitinib (5 μM, Selleckchem) for 4 h, then cultured in the presence or absence of indicated cytokines. After 2 d of cytokine treatments, cells were washed with sterile Dulbecco’s Phosphate Buffered Saline (Gibco) and continued to culture in regular DMEM medium for another two days before measurement of melanin content.

### 2.5 Western blotting

Whole-cell extracts were lysed using Pierce RIPA buffer (Thermo Scientific) or NP-40 buffer supplemented with 1x Halt protease inhibitor cocktail (Thermo Scientific) and 1x Halt phosphatase inhibitor cocktail (Thermo Scientific). The concentration of cell extracts was measured by the Bio-Rad Protein Assay following manufacturer’s protocol. The same amounts of protein extracts were separated on the 4%-20% Mini-Protean TGX gel system (Bio-Rad) and transferred to PVDF (0.45 μm pore size, Millipore) membranes. Membranes were then incubated with the following primary antibodies: Stat1 (D1K9Y, 1:1000, Cell Signaling Technology), pStat1 (Y701, 58D6, 1:1000, Cell Signaling Technology), Stat3 (124H6, 1:1000, Cell Signaling Technology), pStat3 (Y705, D3A7, 1:2000, Cell Signaling Technology), Irf1 (D5E4, 1:1000, Cell Signaling Technology), Tyr (a-PEP7h’, 1:5000), Tyrp1 (a-PEP1, 1:5000), Dct (a-PEP8, 1:5000), Mitf (D5G7V, 1:1000, Cell Signaling Technology), Pmel/gp100 (EP4863(2), 1:1000, Abcam), and Gapdh-HRP (D16H11, 1:1000, Cell Signaling Technology). The secondary antibodies used for detection were HRP-conjugated goat anti-mouse and goat anti-rabbit IgG (1:5000, Thermo Scientific). Band intensities of Tiff images were quantified by using Image J software.

### 2.6 CRISPR-Cas9 Mediated Knockout of *Stat1* and *Stat3*

We designed two guide RNAs targeting different exons of Stat1 and Stat3 loci by online CRISPR Design Tool. Plasmid construction and molecular cloning were done by following the previously published protocol (Ran et al., 2013).

The Cas9 expression construct pSpCas9(BB)-2A-GFP was purchased from Addgene (Plasmid ID 44758). Stat1 (NM_001205313.1) and Stat3 (NM_213659) were used to search gRNA using the online CRISPR Design Tool (http://tools.genome-engineering.org) (Ran et al., 2013).Two different gRNA targeting different exons were used for both Stat1 and Stat3. The sequence of gRNA_Stat1_#1: GGAAACTGTCATCGTACAGC. The sequence of gRNA_Stat1_#2: GGTCGCAAACGAGACATCAT. The sequence of gRNA_Stat3_#1: GCAGCTGGACACACGCTACC. The sequence of gRNA_Stat3_#2: TTCTTCACTAAGCCGCCAAT. Plasmid construction and molecular cloning were done by following the previously published protocol (Ran et al., 2013). B16N cells were transfected with each constructs using Lipofectamine 3000 (Invitrogen) following the manufacturer’s protocol. Single GFP+ cell was sorted into each well of multiple 96-well plates by BD Influx Cell Sorter after 48 h post-transfection. Selected clones were screened for expression of either Stat1 or Stat3 by quantitative real-time PCR, Western blot analysis, and Surveyor mutation detection assay.

### 2.7 Quantitative Real-time PCR

Cells were lysed by Trizol reagent (Invitrogen) and RNA was purified by RNeasy Mini Kit with DNase I digestion (Qiagen) following the manufuatuer’s protocol. Generation of cDNA was performed by GoTaq 2-step RT system (Qiagen). Real-time PCR reactions were measured by ABI StepOnePlus system using SYBR green qPCR master mix (ThermoFisher). The sequence of primers for amplification of different genes were: *Mitf-M* (Forward 5’-GCCTTGTTTATGGTGCCTTC -3’, Reverse 5’GTCCTCCTCCCTCTACTTTCTGT-3’); *Dct* (Forward 5’ CTTCCTAACCGCAGAGCAAC-3’, Reverse 5’ CAGGTAGGAGCATGCTAGGC-3’); *Tyrp1* (Forward 5’ TCTCTTCGGGCAATTAACAG-3’, Reverse: 5’-GGGGAGGACGTTGTAAGATT -3’); *18s rRNA* (Forward 5’-CTTAGAGGGACAAGTGGCG-3’, Reverse 5’-ACGCTGAGCCAGTCAGTGTA-3’). *18s rRNA* was used as the reference control. The ΔΔCT method was used to calculate relative expression level.

### 2.8 Tyrosinase Activity Assay

The tyrosinase activity was determined by measuring the rate of oxidation of 3, 4-Dihydroxy-L-phenylalanine (L-DOPA) as previously described (Newton et al., 2007) with some modifications. B16N were collected and washed with ice-cold DPBS after treatment of indicated cytokines for 4 d. The cell pellets were lysed in 0.1 M phosphate buffer (pH 6.8) containing 1% Triton X-100 and 1x Halt Protease inhibitor cocktail. The lysates were centrifuged at 13,000xg at 4º C for 10 min. 100 μL technical triplicates of tyrosinase-containing supernatants were incubated with 100 μl of 3 mg/mL L-DOPA solutions (Sigma) that were dissolved in 0.1 M phosphate buffer (pH 6.8) at 37º C for 2 h. The 490 nm absorbance was measured at 0 h and 2 h using a multi-well spectrophotometer. The tyrosinase activity was calculated as Δ OD_490_ (OD_490_^2h^-OD_490_^0h^)/min/mg of protein lysate used.

### 2.9 Glycosidase Digestion

B16N cells were collected after culturing for 4 d in the presence or absence of indicated cytokines. 20 μg of protein lysates of each condition was treated with either Endo H (New England Biolabs) or PNGase F for 1 h at 37º C, following the manufacturer’s protocol. After the digestion, the same amount of protein digestion products was subjected to Western blotting analysis probing with PEP7h antibody (1:5000) or Gapdh-HRP (D16H11, 1:5000, Cell Signaling Technology).

### 2.10 Cycloheximide Pulse-Chase Assay

After 4 d treatment of indicated cytokines, B16N cells were washed with DPBS, followed by culturing in medium containing 50 μM of cycloheximide (Sigma) for indicated time points. Cells were collected at 0 h, 3 h, and 6 post cycloheximide treatment. Protein lysates were collected and analyzed for the tyrosinase abundance by Western blotting described before. The tyrosinase degradation rates were estimated by linear regression analysis by GraphPad Prism software.

### 2.11 Statistical Analysis

All bar graphs were generated by GraphPad Prism. Two-tailed unpaired Student’s t-test was used to determine the statistical difference between two groups. For the tyrosinase degradation rates, an analysis of covariance (ANCOVA) was used to determine whether the slopes and intercepts are significantly different. P<0.05 was considered as statistically significant.

## 3. RESULTS

### 3.1 IFNG treatment induces melanogenesis in melanoma cells

To evaluate the effect of interferon cytokines on melanin synthesis, B16 and B16N mouse melanoma cell lines were cultured continuously in medium containing either type I interferon (Ifna2), type II interferon (Ifng), or αMSH as a positive control for the indicated time periods (Figure 1a-d). Treatment of B16 and B16N cells with Ifng for a short period of time (less than 3 d) did not visibly increase melanin synthesis that could be visualized in the cell pellets (Figure 1a, c) or measurable by the melanin content assay (Figure 1b, d). As expected, treatment with αMSH elicited a rapid increase in melanin synthesis, as early as 1 d. Intriguingly, melanin content of B16 and B16N cells was significantly increased after the treatment with Ifng for a longer period of time (at least 3 d) (Figure 1a-d). However, type I interferon Ifna2 failed to affect the melanin content in these two cell lines at any time point (Figure 1a-d). In addition, recombinant human IFNG also increased melanin synthesis in human melanoma cell line Hs 936.T after prolonged treatment, but not after short treatment (Figure 1e, f). Surprisingly, αMSH inhibited melanogenesis in this cell line (Figure 1e, f).

**Figure 1.**
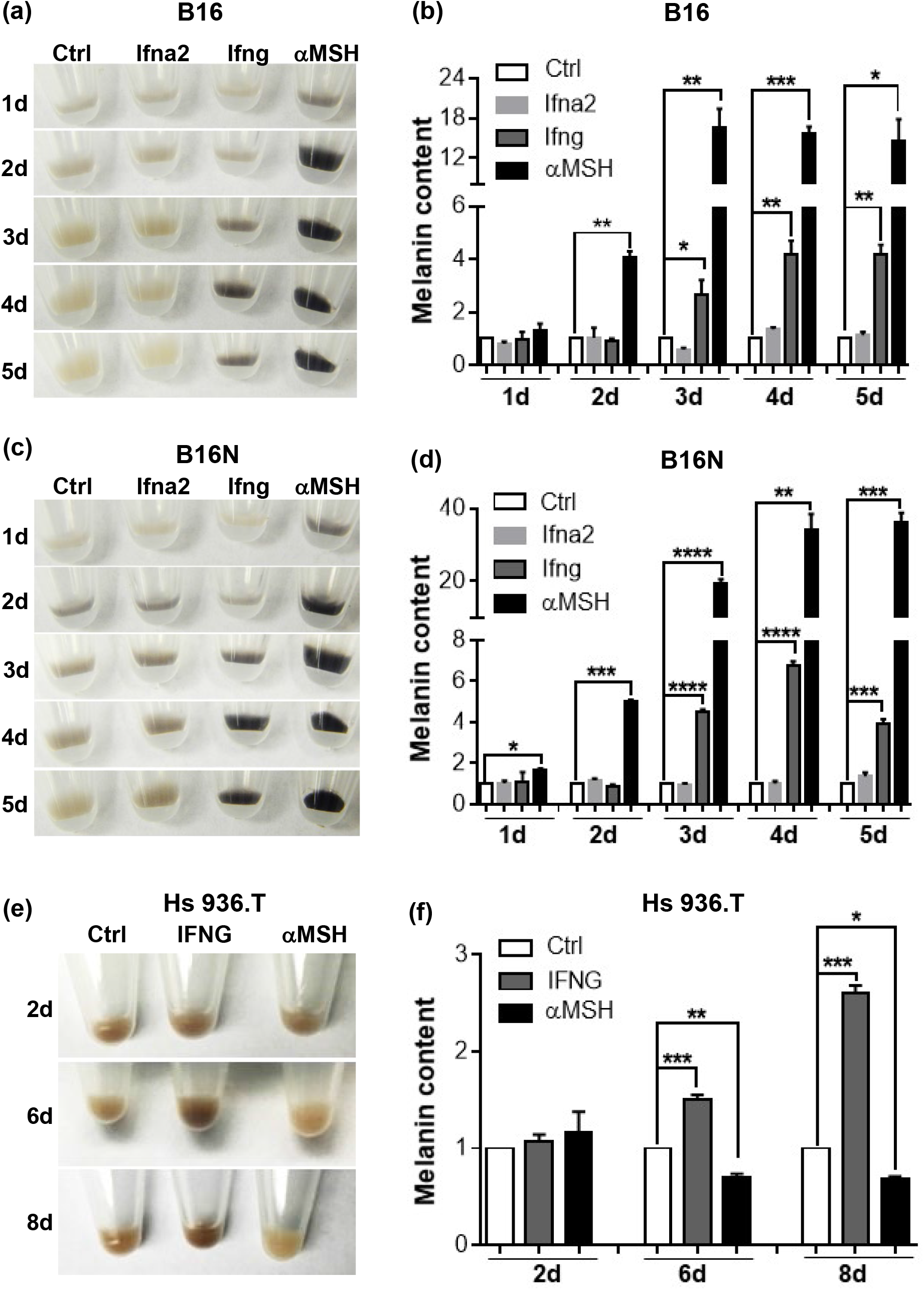
Effects of cytokines on melanin synthesis. (a), (c), (e) Images of cell pellets of B16, B16N, and Hs 936.T, respectively, following cytokine treatments for indicated time points. (b), (d), (f) Quantification of melanin content in melanoma cells in the presence or absence of cytokines treatments for indicated time points. Images are representative of at least 3 independent experiments. All graphed data are presented as the mean ± SEM of three biological replicates, relative to the Ctrl group of each indicated time points. *P<0.05; **P<0.01; ***P<0.001; ****P<0.0001.

To test if prolonged and continuous Ifng signaling is necessary for inducing melanin synthesis, we discontinued Ifng treatment at the indicated time points, as low as 1 h, but continued to culture the B16 and B16N cells in regular media, followed by measurement of the melanin content at 4 d (Figure 2). Unexpectedly, a visible elevation in melanin content was apparent in melanoma cells by exposure to Ifng as short as 1 d (Figure 2), which demonstrated that Ifng-induced melanogenesis did not require continuous treatment.

**Figure 2.**
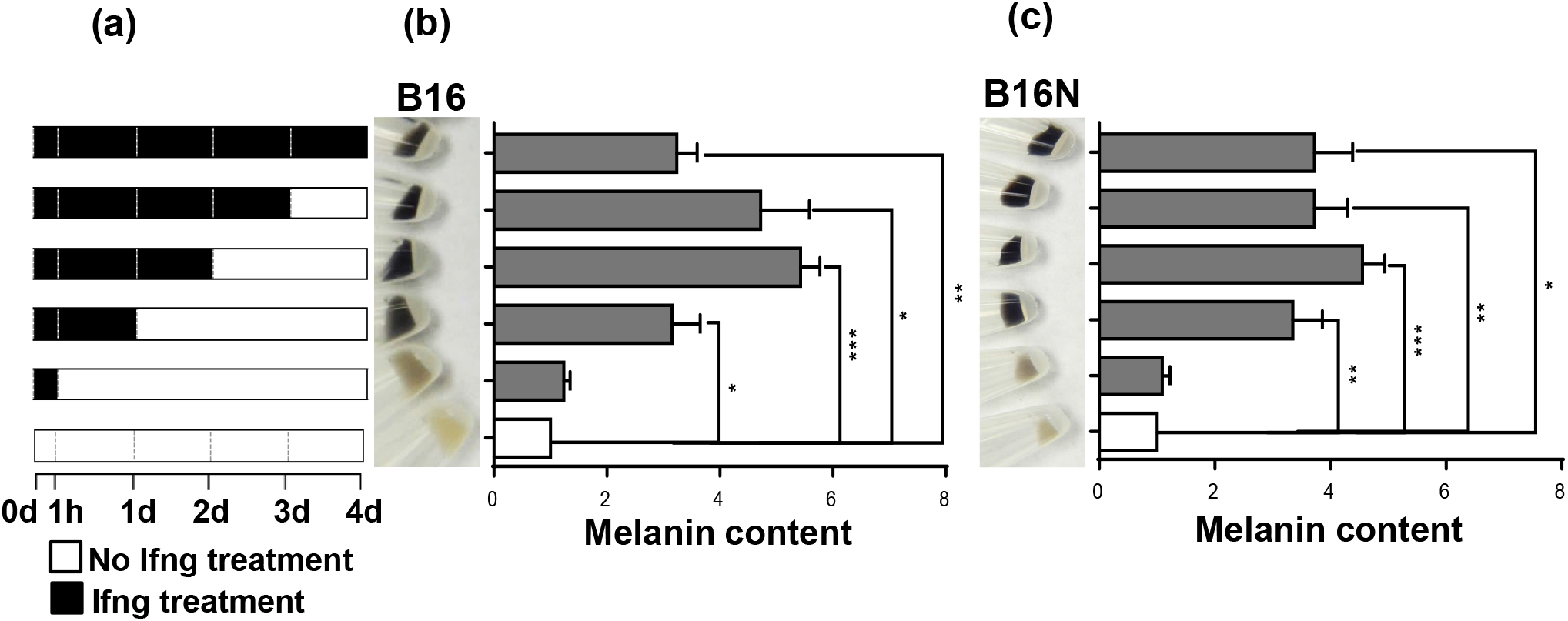
Temporal effects of Ifng treatment on melanogenesis. (a) Schematic of Ifng treatment schedule. Cells were collected after 4 d of culture in the presence or absence of Ifng for indicated time points. (b), (c) Image of cell pellets collected after 4 d of treatment and melanin content quantification in B16 and B16N cells, respectively. The graphed data are presented as the mean ± SEM of three biological replicates, relative to the Ctrl group of each indicated time points. *P<0.05; **P<0.01; ***P<0.001.

### 3.2 JAK-STAT1 mediate IFNG-induced melanogenesis

The JAK-STAT cascade is the major mediator of IFNG signaling to perform its downstream biological effects. However, upon IFNG binding to its receptors, several other pathways, such as MAP kinase and PI3K pathways, are activated independently in addition to the canonical JAK-STAT axis to maximize the biological functions of IFNG (Gough et al., 2008). To test the requirement of the Jak1/2 route for Ifng-induced melanin synthesis, we pretreated the cells with the potent Jak1/2 inhibitor ruxolitinib. Ruxolitinib (5 μM) pre-treatment completely abolished Ifng-induced melanogenesis in B16N cells (Figure 3a). In contrast, ruxolitinib treatment did not affect the basal melanin level or αMSH-induced melanogenesis (Figure 3a). Surprisingly, we observed a significant synergistic effect of Ifng and αMSH in enhancing melanogenesis (Figure 3a).

**Figure 3.**
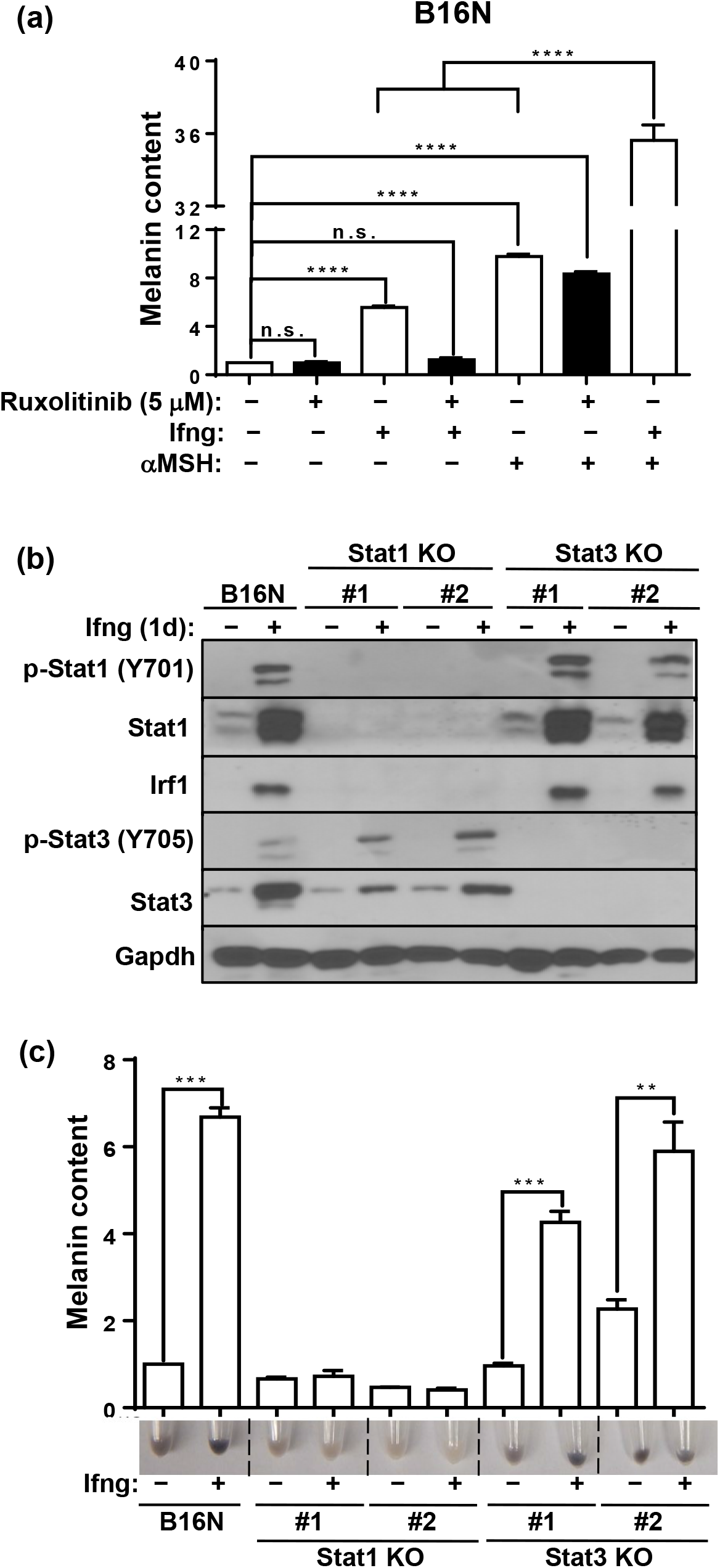
Jak1/2-Stat1 mediates Ifng-induced melanogenesis. (a) Melanin content quantification in B16N cells cultured with indicated cytokines in the presence or absence of ruxolitinib (5 μM). B16N cells were pretreated with ruxolitinib for 1 h followed treated with or without indicated cytokines for 2 d and collected at 4 d. (b) Western blot of pStat1 (Y701), Stat1, Irf1, pStat3 (Y705), Stat3, and Gapdh expressions in parental B16N cells and Stat-KO clones in response to Ifng treatment. (c) Image of pellets and melanin content quantification of B16N and Stat-KO clones collected after 4 d of treatment with or without Ifng. Immunoblotting images are representative of two independent experiments in (b). Data are presented as mean ± SEM of three biological replicates in (a) and (c). **P<0.01; ***P<0.01; ****P<0.0001.

STAT1 is the predominant downstream transcription factor that mediates IFNG signaling. Accumulating evidence suggests that STAT3 is also activated directly upon IFNG treatment (Qing and Stark, 2004; Zaidi and Merlino, 2011) and its activity is important in maintaining melanogenesis in B16 cells (Yang et al., 2010). Indeed, We observed that both pStat1 (Y701) and pStat3 (Y705) were robustly activated in response to Ifng treatment (Figure 3b). To evaluate which transcriptional factor mediates the Ifng-induced melanogenesis, we generated individual Stat1- or Stat3-knockout (KO) clones of B16N cells by CRISPR-Cas9 gene-editing. The upregulation of the transcription factor Irf1 upon Ifng treatment was completely blocked in the Stat1-KO clones; while its activation in Stat3-KO clones remained intact (Figure 3b). Interestingly, Ifng treatment increased Stat3 protein level, which was partially inhibited by loss of Stat1, which suggests that the upregulation of Stat3 is at least partially dependent on Stat1 (Figure 3b). In contrast, activation of pStat1 by Ifng was unaffected in Stat3-KO cells (Figure 3b). Melanin content assay showed that Ifng failed to increase melanin synthesis in Stat1-KO clones but not in Stat3-KO clones (Figure 3c). These data provide evidence that Ifng-induced melanogenesis is mediated by the Jak1/2-Stat1 signaling axis.

### 3.3 Transcriptional and translational effects of IFNG on melanogenesis-associated gene expression

The MITF-M isoform of MITF is exclusively expressed in the melanocyte compartment and acts as a master regulator of melanogenesis (Wang et al., 2010). In contrast to the rapid upregulation of Mitf-M mRNA by αMSH as expected, Ifng and Ifna2 decreased the mRNA of Mitf-M after 2 h treatment of B16N cells (Figure 4a). Mitf is known to regulate the transcription of melanogenic enzymes including tyrosinase (Tyr), tyrosinase-related protein 1 (Tyrp1), and dopachrome tautomerase (Dct) (Kawakami and Fisher, 2017; Sturm, 2006). However, despite the increasing Mitf expression after treatment with Ifng for 1 d, the mRNA expression levels of Tyr, Tyrp1, and Dct were not upregulated by either short or prolonged treatment with Ifng (Figure 4b, c, d). Protein levels of Tyr, Tyrp1, and Dct were unchanged by treatment of Ifng for 1 d (Figure 4e, f), and melanin production did not increase during this time frame. However, at the protein level, Tyr was significantly increased at 4 d by treatment of Ifng; while the protein levels of Tyrp1, Dct, and Pmel remained unaffected (Figure 4g, h). The significantly increased Tyr protein level was in accordance with the increasing melanin content. We confirmed this by measuring tyrosinase activity, which is a more sensitive indicator of melanogenesis than measurement of total melanin content (Newton et al., 2007). Prolonged treatment of B16N cells with Ifng profoundly increased Tyrosinase activity (Figure 5a), which was closely correlated with the melanin content and Tyr protein level. In addition, we found that the increase in Tyr protein was dependent on Stat1, as the Stat1-KO clones failed to show increased Tyr in response to Ifng treatment (Figure 5b). As expected, Ifna2 had no effects on the expression of melanogenesis-associated gene expression, including Tyr activity (Figure 4a-h and Figure 5a). These data suggest that Ifng/Stat1-induced melanogenesis is not dependent on the upregulation of Mitf or mRNA level of Tyr; rather, it is dependent on the increase in the Tyr protein abundance.

**Figure 4.**
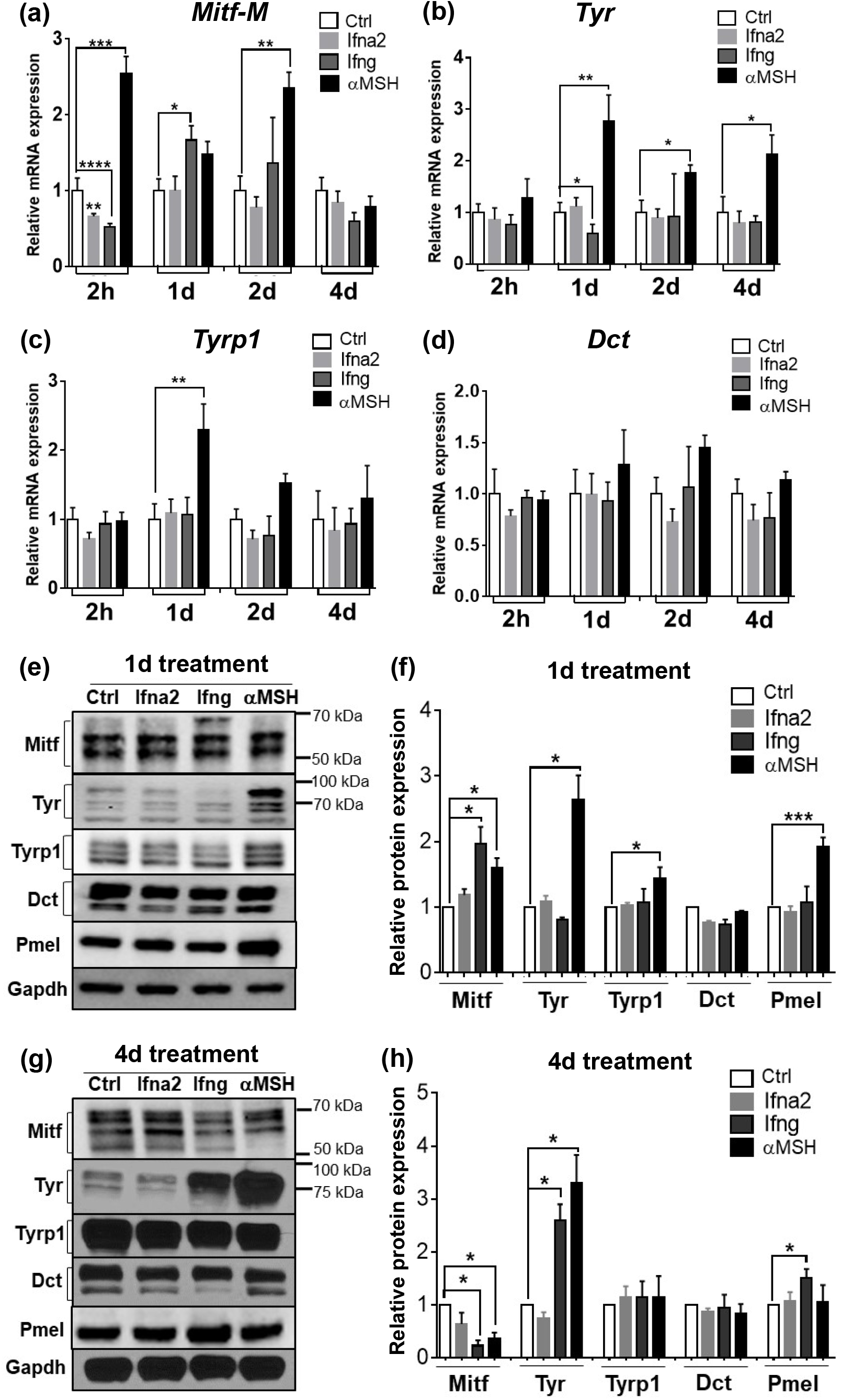
Effects of cytokines on the expression of melanogenesis-related genes. qRT-PCR analysis of mRNA expression of *Mitf-M* (a), *Tyr* (b), *Tyrp1*(c), and *Dct* (d) in B16N upon cytokine treatments for indicated time points. Bargraph is presented as mean ± SEM of three biological replicates. (e) Western blot of Mitf, Tyr, Tyrp1, Dct, and Pmel in B16N cells upon indicated cytokine treatment for 1 d (e) and 4 d (g). (f), (h) Quantification of protein abundances of Western blot. Gapdh was used as a loading control. The ratio of target protein/Gapdh is relative to the Ctrl group. Data are presented as mean ± SEM of 3-5 independent experiments. *P<0.05; **P<0.01; ***P<0.001; ****P<0.0001.

**Figure 5.**
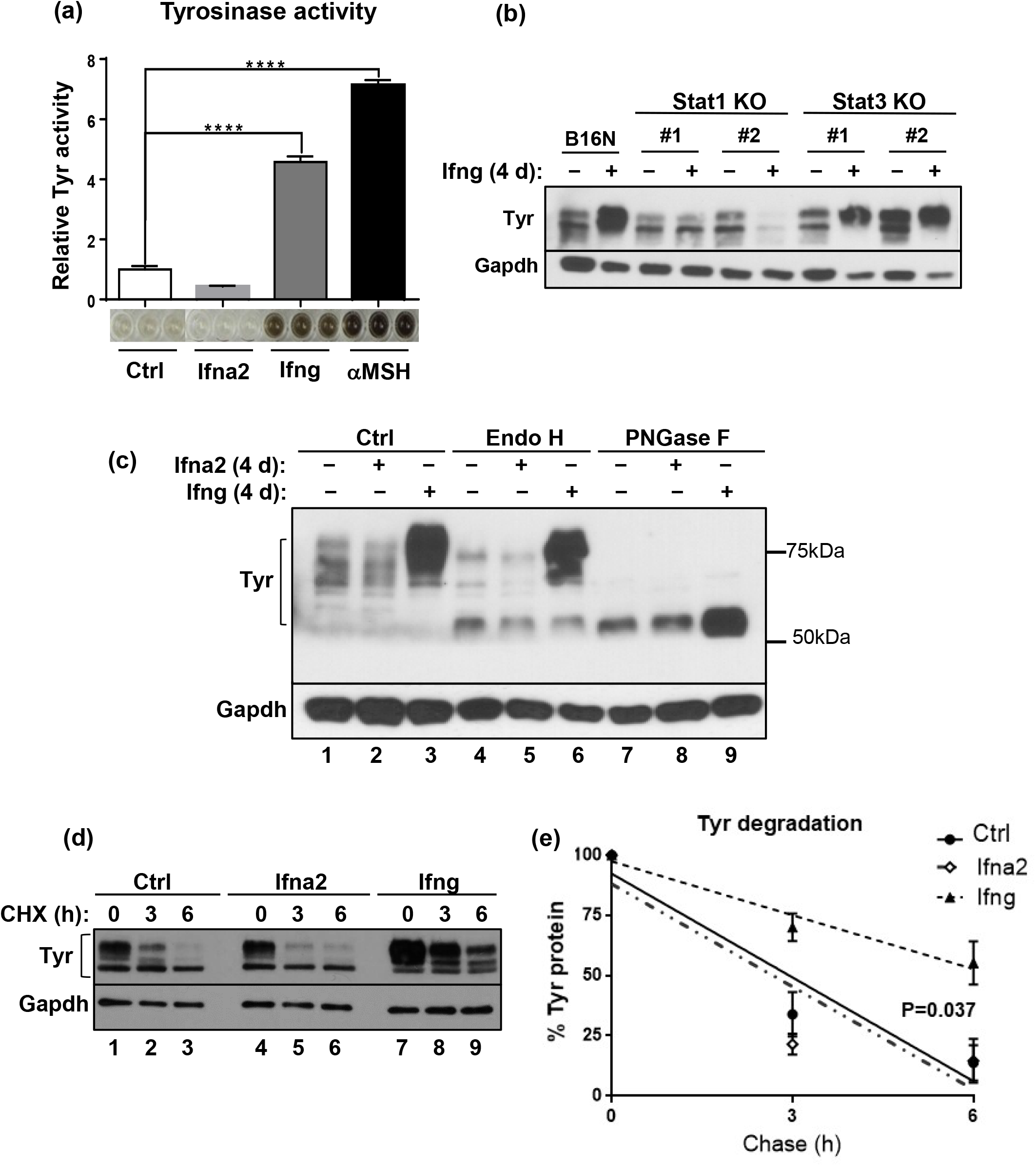
Accumulation of ER-resistant, fully mature, and prolonged half-life of tyrosinase in response to Ifng treatment. (a) Tyrosinase activity of B16N cells treated with different cytokines. The histogram is presented as mean ± SEM of three biological replicates, relative to the Ctrl group. (b) Western blot of Tyr expression in B16N and Stat-KO cells treated with or without Ifng for 4 d. The Western blot is representative of two independent experiments. (c) Cell lysates obtained from B16N cells following 3 d of treatment with either Ifna2 and Ifng were subsequently incubated with the glycosidases, Endo H and PNGase F. Western blot of Tyr to confirm the extent of digestion. Tyr is range from 58 to 80 kDa. (d) B16N cells were cultured with either Ifna2 or Ifng for 4 d, then treated with 50 uM cycloheximde for 3 h or 6 h, in following of Western blot of Tyr abundance. (e) The curve represents the mean ± SEM of three independent measurements of Western blot of (d). Y-axis, the percentage of Tyr/Gapdh is relative to Ctrl 0 h. Linear regression and ANCOVA were performed in Prism 6. *P<0.05; ****P<0.0001.

### 3.4 IFNG treatment causes prolonged half-life and accumulation of Tyrosinase

Western blotting for Tyr showed that Ifng treatment for 4 d led to a significant increase in the abundance of mature Tyr that appeared mostly as a ∼75-80 KDa band (Figure 5c, lanes 1-3). We utilized glycosidase digestion to evaluate the post-translational modification of Tyr following cytokine treatments. Endoglycosidase H (Endo H) cleaves early, less processed mannose added to Tyr in the ER (Endo H-sensitive), but not from the mature Tyr that exits the TGN (Endo H-resistant) (Ando et al., 2006). Another enzyme used for glycosidase digestion, N-Glycosidase F (PNGase F) cleaves all N-linked oligosaccharides from glycoproteins. PNGase F digestion showed that the total amount of deglycosylated Tyr (∼58kD) significantly increased after treatment with Ifng but not by Ifna2 (Figure 5c, lanes 7-9). The Endo H digestion revealed that the majority of Tyr extracted from Ifng-treated B16N cells was the Endo H-resistant mature form (Figure 5c, ∼75kD band in lane 6); whereas, Tyr in either untreated or Ifna2-treated B16N cells was largely Endo H-sensitive immature protein (Figure 5c, ∼58kD bands in lanes 4 and 5, respectively.)

To investigate whether Ifng-mediated glycosylation promotes Tyr protein stability and an increased half-life, we performed cycloheximide pulse-chase assay. The results showed that the half-life of Tyr in untreated cells was approximately 3 h (Figure 5d, lanes 1-3 and Figure 5e), as reported earlier (Ando et al., 2004; Halaban et al., 1997); but interestingly, the half-life of Tyr was significantly prolonged in Ifng-treated cells (Figure 5d, lanes 7-9, and Figure 5e). In contrast, Ifna2 did not alter the degradation rate of Tyr (Figure 5d, lanes 4-6, and Figure 5e). Intriguingly, there was a reduced amount of ER-sensitive Tyr in Ifng-treated B16N, suggesting that it did not increase de novo synthesis or early processing of Tyr in the ER, which was consistent with the mRNA expression of Tyr. Therefore, our findings suggest that Ifng leads to the accumulation of the mature form of Tyr with a prolonged half-life, results in increased melanin synthesis.

## 4. Discussion

Two previous studies had shown that IFNG inhibited melanogenesis leading to hypopigmentation in melanocytes and B16 melanoma cells (Natarajan et al., 2014; Son et al., 2014). Natarajan et al. showed that IFNG signaling caused hypopigmentation in human primary melanocytes by arresting melanosome maturation. This effect was accompanied by reduced expression of the melanogenic genes TYR, DCT, and TRP1, but was interpreted to be independent of MITF (Natarajan et al., 2014). In contrast, Son et al. reported that IFNG inhibited melanogenesis by suppressing the expression of Mitf, which led to reduced activation of the *Tyr* promoter in B16 cells (Son et al., 2014). Here we have identified an entirely new regulatory mechanism of IFNG-mediated enhancement of melanogenesis that is independent of Mitf and is based on the post-translational modification (glycosylation) of Tyr. These opposing results can only be explained in terms of the extremely intricate genetic control of melanogenesis that would be manifest in the remarkable heterogeneity of the pigmentary phenotype observed in the human primary melanocytes obtained from different individuals and mouse melanoma cells. Nevertheless, it seems plausible that IFNG is capable of suppressing as well as enhancing melanogenesis in the melanocytic compartment in a genetic and/or cellular context-dependent manner. Elucidation of these differing contexts will require further work.

Natarajan et al. also showed that *Ifng*^*−/−*^ mice in the C57BL/6 strain background exhibit a hyperpigmented tail phenotype and that pigmentation is further enhanced by exposure to UVB radiation (Natarajan et al., 2014). We have not observed this phenotype in *Ifng*^*-/-*^ mice as compared to wildtype littermates. In fact, we have determined that *Ifng*^*-/-*^ mice have reduced melanocytic activation and migration to the epidermis in response to UVB irradiation (data not shown), which seems to make a UVB-induced hyperpigmentation phenotype in *Ifng*^*-/-*^ mice implausible. It is also important to note that transgenic mice in which the AU-rich element (ARE) in the 3’UTR of the *Ifng* gene is deleted (*Ifng-ARE-Del*), with the homozygous *ARE-Del*^*-/-*^ mice exhibiting chronic production of circulatory Ifng, exhibit visibly elevated skin pigmentation (Bae et al., 2016; Hodge et al., 2014), which supports a melanogenesis-inducing effect of Ifng signaling in melanocytes.

Numerous studies have shown the role of Mitf in initiating and maintaining the transcription of Tyr, Tyrp1, and Dct in melanocytes and melanoma cells. Unexpectedly, Ifng did not increase the mRNA expression of Tyr, Tyrp1, and Dct despite an increase in Mitf expression. In contrast, upregulation of Mitf by αMSH elevated mRNA expression of Tyr and Tyrp1. One possible explanation for these results is that Ifng induces factors that cooperate with Mitf to negatively regulate the transcription of *Tyr* and *Tyrp1*. It has been reported that Mitf-induced transcription of Tyr, Tyrp1, and Dct is dependent on other transcriptional factors, such as Sox10 (D’Mello et al., 2016; Jager et al., 2006; Potterf et al., 2001). We observed a decrease in the expression of Sox10 by Ifng treatment (data not shown), which might explain why mRNA expression of Tyr was unresponsive to the upregulation of Mitf. The enhanced stability and consequent accumulation of postranslationally modified Tyr protein can readily explain the lag in the induction of melanin biosynthesis in response to Ifng as compared to αMSH treatment. Newton et al. have also reported that the elevation of MITF expression by forskolin induces tyrosinase protein expression, but not its mRNA expression (Bae et al., 2016; Newton et al., 2007).

Interestingly, Yang et al. have proposed that the constitutive activation of Stat3 in B16 cells was important for tyrosinase expression (Yang et al., 2010). They mutated the phosphorylation site of Stat3 at Y705, which resulted in the loss of its transcriptional activity. The overexpression of mutated dominant-negative Stat3 (Stat3-DN) in B16 cells decreased Tyr expression, which led to the inhibition of melanogenesis (Yang et al., 2010). However, we have not observed a reduction in Tyr expression and melanogenesis in Stat3-KO clones. Additionally, we have shown that Ifng-induced hyperpigmentation was mediated by the Stat1, but not the Stat3, signaling axis. These results suggest that the depigmentation effect mediated by Stat3-DN is not due to the disruption of the transcriptional function of Stat3. Further work is needed to elucidate the exact mechanism.

IFNG has long been known as the principal proinflammatory cytokine that plays the role of the central orchestrator of inflammation and autoimmune disease (Schroder et al., 2004). IFNG is predominantly produced by the NK, NKT, and macrophages of the innate immune system, as well as the CD4+ Th1 and CD8+ cytotoxic T lymphocytes of the adaptive immune system (Schroder et al., 2004; Zaidi, 2019; Zaidi and Merlino, 2011). IFNG is directly involved in almost all types of skin inflammatory conditions, e.g. those caused by pathogenic infections, tissue injury, tissue stress, and/or malfunction (Dries and Perry, 2002; Xiao et al., 2009). One of the most prevalent skin conditions is post-inflammatory hyperpigmentation (PIH), which is characterized by a localized increase in melanogenesis by melanocytes (hypermelanosis) following numerous skin inflammatory conditions. Some of the most common inflammatory conditions that cause PIH are acne, atopic dermatitis, allergic contact dermatitis, and several types of dermatological procedures (Kaufman et al., 2018). Wound healing following skin abrasions, cuts, and burns also frequently lead to PIH. Exposure to the ultraviolet radiation (UVR) from the sun further exacerbates PIH (Ruiz-Maldonado and Orozco-Covarrubias, 1997). Overall, PIH is one of the most common skin disorders and affects hundreds of millions of people worldwide (Park et al., 2016). At the histopathological level, PIH features elevated deposition of melanin in either the epidermis or the dermis. Both patterns are due to increased melanogenic activity without an increase in the number of melanocytes (Park et al., 2017). The molecular mechanisms of the induction of PIH remain poorly understood. In light of our results presented here, it is plausible to speculate that inflammation-associated release of IFNG in the skin microenvironment is capable of inducing melanogenesis in the inflamed skin, leading to PIH. Elucidating this potential causal connection is an enticing future prospective investigative avenue.

